# Interplay Between Intrinsically Disordered Proteins and Atomically Precise Gold Nanoclusters Modulates their Optical Properties

**DOI:** 10.1101/2025.05.06.652423

**Authors:** Santiago Rodriguez, Sylvain Kumanski, Zeineb Ayed, Aurélie Fournet, Charlène Bouanchaud, Amin Sagar, Frédéric Allemand, Ute Resch-Genger, Juan Cortés, Nathalie Sibille, Fabien Chirot, K. David Wegner, Rodolphe Antoine, Xavier Le Guével, Pau Bernadó

**Affiliations:** Centre de Biologie Structurale, Université de Montpellier, INSERM, CNRS, 29 rue de Navacelles, 34090, Montpellier, France; Université Grenoble Alpes, INSERM U1209, CNRS UMR 5309, Institut pour l’Avancée des Biosciences (IAB), Cancer Targets & Experimental Therapeutics, 38000 GRENOBLE, France; Institut Lumière Matière, UMR5306, Université Claude Bernard Lyon1-CNRS, Université de Lyon, 69622, Villeurbanne Cedex, France; BicycleTx Limited, Cambridge, UK; Federal Institute for Materials Research and Testing (BAM), Division Biophotonics, Richard-Willstaetter-Str. 11, 12489 Berlin, Germany; LAAS-CNRS, Université de Toulouse, CNRS, F-31400 Toulouse, France

**Keywords:** Gold nanoclusters, intrinsically disordered proteins, Small-Angle X-ray Scattering (SAXS), photoluminescence, NIR-II

## Abstract

Understanding how structural and optical properties of metallic nanoclusters can be tuned by proteins is crucial for the use of these hybrid molecules in biomedical applications. The interaction of proteins with ultrasmall, atomically-precise gold nanoclusters (Au-NCs) has been mainly investigated in the context of structured proteins, while their behavior with intrinsically disordered proteins (IDPs) remains unexplored. This work examines the structural and optical properties of Au-NCs interacting with bioengineered IDPs containing up to three cysteines. We show that, by exploiting the conformational flexibility of cysteine-containing IDPs, we can anchor proteins to Au-NCs in a position-specific manner, leading to new bioconjugates with properties that differ from those of the individual components. We observed an up to 15-fold photoluminescence enhancement depending on the number of cysteines anchored. By combining mass spectrometry, small-angle X-ray scattering (SAXS), and computational modelling, the ensemble structures of nine bioconjugates with different stoichiometries were elucidated, indicating their overall compactness. Our results suggest that the interface between these atomically-precise species and the conformationally fluctuating protein is responsible for the optical properties of these nanobioconjugates. This research improves our understanding of Au-NC– protein interactions, paving the way to novel nano-molecular hybrid conjugates with tunable properties for bioimaging and therapeutic applications.

## Introduction

The design of hybrid nanobioconjugates has attracted increasing interest, driven by the need for innovative platforms in sensing, imaging and therapeutics^1,2^. A particularly promising avenue lies in the synergistic development of atomically precise metal nanoclusters—primarily gold nanoclusters (Au-NCs)—anchored to proteins, leading to a new generation of functional nanobioconjugates^3–6^. These hybrid systems leverage the unique physico-chemical and optical properties of Au-NCs, including fluorescence, catalytic activity, chirality, and photo- or radio-responsiveness, while integrating the biological functions and properties of proteins.

Despite significant progress in synthesizing and characterizing these constructs, a fundamental understanding of the Au-NC-protein interface and its role in tuning the characteristics of nanoclusters remain incomplete^7^. Recent experimental studies, complemented with time-dependent density functional theory (TDDFT) and molecular dynamics simulations of Au-NCs covalently or electrostatically bound to the Bovine Serum Albumin (BSA), have provided some hints on the relevance of the interface^8,9^. These studies suggest that key parameters, such as protein:Au-NC stoichiometry, binding nature and environment, strongly influence the optical and electronic properties of these nanobioconjugates. Moreover, the intrinsic characteristics of proteins, such as their conformation and electrostatics, likely play a crucial role in modulating the overall stability, optical properties, and functionality of these hybrid systems. A deeper exploration of the latter factors is essential for unlocking the full potential of Au-NC–protein conjugates for advanced biomedical applications. While most studies on the interaction of Au-NCs and proteins have focused on globular proteins^5^, Michael Assfalg and coworkers pioneered the study of ultrasmall gold nanoparticles and a fragment of protein tau, an IDP involved in Alzheimer’s disease^10^. They found that the nanoparticles serve as a seed for the formation of stable non-covalent multimolecular assemblies, showing a new type of nano–bio interaction^10^.

## Results

In the present study, we investigate the ability of engineered IDPs with different number of free cysteines to modulate the properties of Au-NC. We first explored the reaction landscape of the atomically precise Au-NC, Au_25_pMBA_18_ (pMBA : para-mercaptobenzoic acid), as a model, in the presence of small IDPs. Precisely, we used the C-terminal disordered tail of the β2-adrenergic receptor (β2), which has two free cysteines in its wild type (β2-2cys), and two single point mutations yielding constructs with 1 (β2-1cys) and 3 (β2-3cys) free cysteines (see Table S1 and methods for details). When mixing the Au-NC with each of the three constructs in a ratio 1:2 (protein:Au-NC) for 3h, a reduced number of species, which differ from the free Au-NC and the proteins, could be identified in the native PAGE-gel in the NIR-II (1000-1700 nm) and also in the red (667nm) region (Fig. S1). In the presence of β2-2cys, the mixture yielded three bands as revealed by fluorescence and Coomassie staining, indicating that the resulting species contained both the Au-NC and the protein. Interestingly, an equivalent number of conjugated species was observed when the number of free cysteines of the protein were one or three. In all cases, the two lower bands closely matched in the gel, while the higher ones migrated similarly as the free proteins. It is interesting to note that the positions of these three bands were different depending on the β2 construct used. This suggests that the molecular weight, size and/or net charge of the bioconjugates changed depending on the number of cysteines.

Then, we explored how the stoichiometry, from 1:0.5 to 1:3, and the reaction times, from 30 minutes to 24h, influenced the population of the resulting species (Fig. S2). For the three β2 constructs, the three main bands were present under the vast majority of experimental conditions. However, at high protein:Au-NC ratios (1:0.5 and 1:1), a fourth band with a migration behavior similar to the free protein appeared concomitantly to the decrease in the population of the lowest bands. Interestingly, at the smallest ratios of 1:2 and 1:3 and short reaction times, the highest band was the major species, but its population slowly decreased with time to enhance that of the lowest bands. These observations confirm that the population of the species formed by mixing the IDP and Au-NC could be modulated by stoichiometry and reaction time, suggesting their transient nature and complex transformation pathways. However, they are not strongly influenced by the number of free cysteines, as the three β2 constructs presented the same overall behavior.

Next, the nanobioconjugates resulting from the mixture of Au-NC with the three β2 constructs were chemically, optically, and structurally characterized. Therefore, we performed large scale reactions, and the resulting mixtures were subjected to Size-Exclusion Chromatography (SEC) (Fig. 1a). In line with the PAGE gels, three peaks were observed for each of the β2 constructs (Fig. S3). While the most compact species (Peak 1) was the major product for the mixture of Au-NC with β2-1cys, the major species for β2-2cys and β2-3cys mixtures corresponded to the middle peak (Peak 2). The three peaks resulting for the three constructs were collected, concentrated, and subjected to HPLC-coupled to mass spectrometry (MS) analysis to identify their chemical composition (see Methods section for details). All the experimentally derived molecular weights (MWs) could be explained as a single Au_25_ cluster with a defined number of proteins and pMBA molecules (Table S2). The major species (Peak 1) of the Au-NC:β2-1cys mixture showed a MW that was in excellent agreement with a single protein attached to an Au-NC replacing a single pMBA molecule. Interestingly, when analyzing the MWs of the major species (Peak 2) for the mixtures of Au-NC with β2-2cys and β2-3cys, the values derived were compatible with a single Au-NC:β2 conjugate in which two and three pMBA molecules were removed, respectively. The correlation between the number of free cysteines of the β2 mutants and eliminated pMBA molecules for the major species strongly suggested that the formation of these conjugates followed a substitution mechanism in which the reduced thiols and pMBA molecules were replaced by cysteines. As a consequence, for the β2-2cys and β2-3cys conjugates, the protein can interact with the Au-NCs at multiple anchoring points, exposing long flexible loops and tails to the solvent (Fig. 1b). To better understand the pMBA/cysteine substitution mechanism, we mixed the Au-NC with an engineered β2 construct without cysteines (β2-0cys, Table S1). Interestingly, the only band appearing in the PAGE-gel as revealed by Coomassie staining, corresponded to that of the protein, indicating that no conjugates were formed when no free cysteines were present in the protein (Fig. S4).

**Figure 1.**
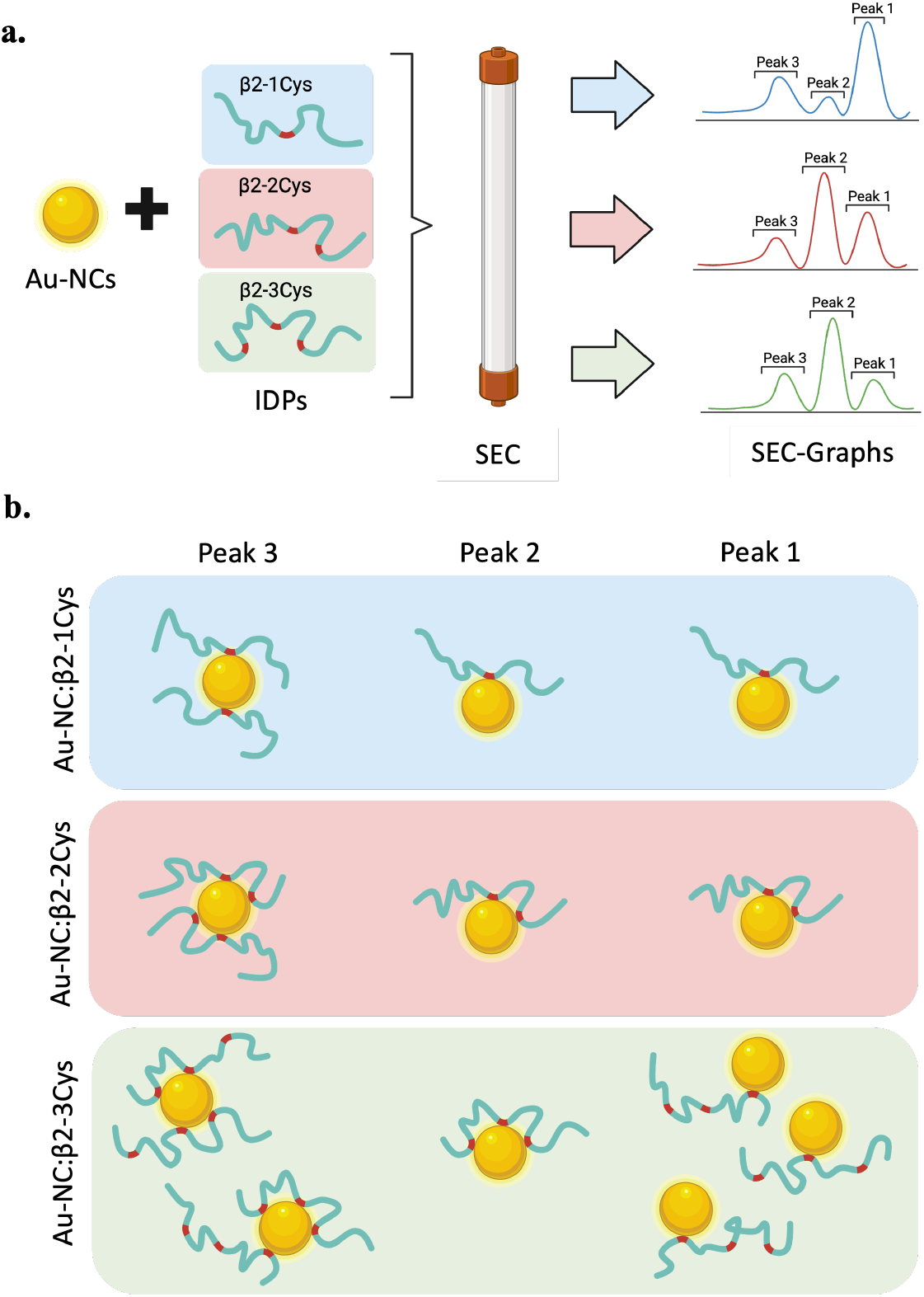
Schematic representation of the protocol followed to synthetize and isolate the species formed upon mixing Au-NCs with the three constructs of the β2 protein (**a**), and their conformation suggested by their mass spectrometry analysis (**b**). Cysteines are highlighted in red within the protein and the Au-NC is represented as a yellow sphere. When multiple arrangements are compatible with the MS data, several cartoons are displayed.

MS data obtained for the minor peaks provided information on the versatility of the Au-NCs to interact with the proteins. In terms of stoichiometry, Peak 2 for Au-NC:β2-1cys and Peak 1 for Au-NC:β2-2cys were indistinguishable from the corresponding major peaks. Interestingly, the MW derived from Peak 1 of Au-NC:β2-3cys indicated that a single pMBA had been removed from the Au-NC, suggesting that this species presented two free cysteines in solution. Given the reactivity of cysteine thiols with Au-NCs, we speculate that the formation of an intramolecular disulfide bond could be responsible for this observation. The MWs derived from the three species with the shortest elution time in the SEC (Peak 3) showed the presence of species with a 1:2 stoichiometry, indicating that these Au-NCs could simultaneously bind two protein chains. Interestingly, the conjugates obtained from β2-1cys and β2-2cys had all the cysteines of both proteins engaged in binding to the Au-NC, implying that up to four pMBA molecules could be removed from the AuNC (Fig. 1). A slightly different scenario occurred for Peak 3 of the β2-3cys conjugate. Even though two proteins forming a complex with the Au-NC, only four out of the six available cysteines were engaged in Au-NC binding. Again, the presence of intra- or intermolecular disulfide bridges hampering the interaction with the gold atoms at the Au-NC surface could not be discarded.

Next, the optical properties of the main species formed by the three β2 constructs with the Au-NCs were investigated. The absorbance spectra of the Au-NCs exhibited characteristic bands and shoulders at 450, 480, 690, and 800 nm, which were attributed to molecular-like transitions occurring within the core (Au_13_ kernel) and shell (Au_n_R_m_ staple motifs) of the Au_25_ nanocluster^11,12^. The complexation of this Au-NC with the three β2 constructs through their cysteines seemed to moderately alter the icosahedral structure of the Au-NC, as confirmed by the slight blue-shift of the band at 690 nm^12^. Note that the increase in absorbance at lower wavelengths was attributed to the absorption of the protein (Fig. S5).

Photoluminescence (PL) spectra of Au-NCs typically exhibit a relatively broad emission band peaking at 1100 nm with a shoulder at 1250 nm. While the exact origin of the NIR-II PL remains unclear, recent studies indicate that the ligand plays a crucial role in modulating the NIR-II fluorescence and phosphorescence through intramolecular charge transfer (ICT)^13,14^. We observed a striking enhancement of the Au-NC PL upon protein conjugation, with the number of cysteines directly influencing the amplification of a shorter-wavelength PL emission at 850 nm (Fig. 2A and S5). Notably, the PL intensity increased more than 15-fold for the complex containing three cysteines with a significant 3-fold increase for two cysteines compared to one cysteine. This enhancement appeared to be wave-length-independent (Fig. S5). PL decay measurements of the Au-NC:β2 complexes revealed a progressive increase in the average intensity weighted lifetime, <τ>_int_, with the number of cysteines, from 133 ns for Au-NC alone to 811 ns for Au-NC:β2-3cys (Fig. 2B and S5). The simultaneous enhancement in PL intensity and average lifetime suggested that the presence of the protein reduces non-radiative recombination processes at the gold surface. Interestingly, this effect is highly dependent on the number and chemical features of the anchoring sites covalently bound to the Au-NCs^8,9^. While a previous study showed a moderate impact on the PL when replacing pMBA by cysteine on atomically precise Au_25_ nanoclusters^8^, our notable PL enhancement suggested a pivotal role of the interface between the disordered protein and the Au-NC.

**Figure 2.**
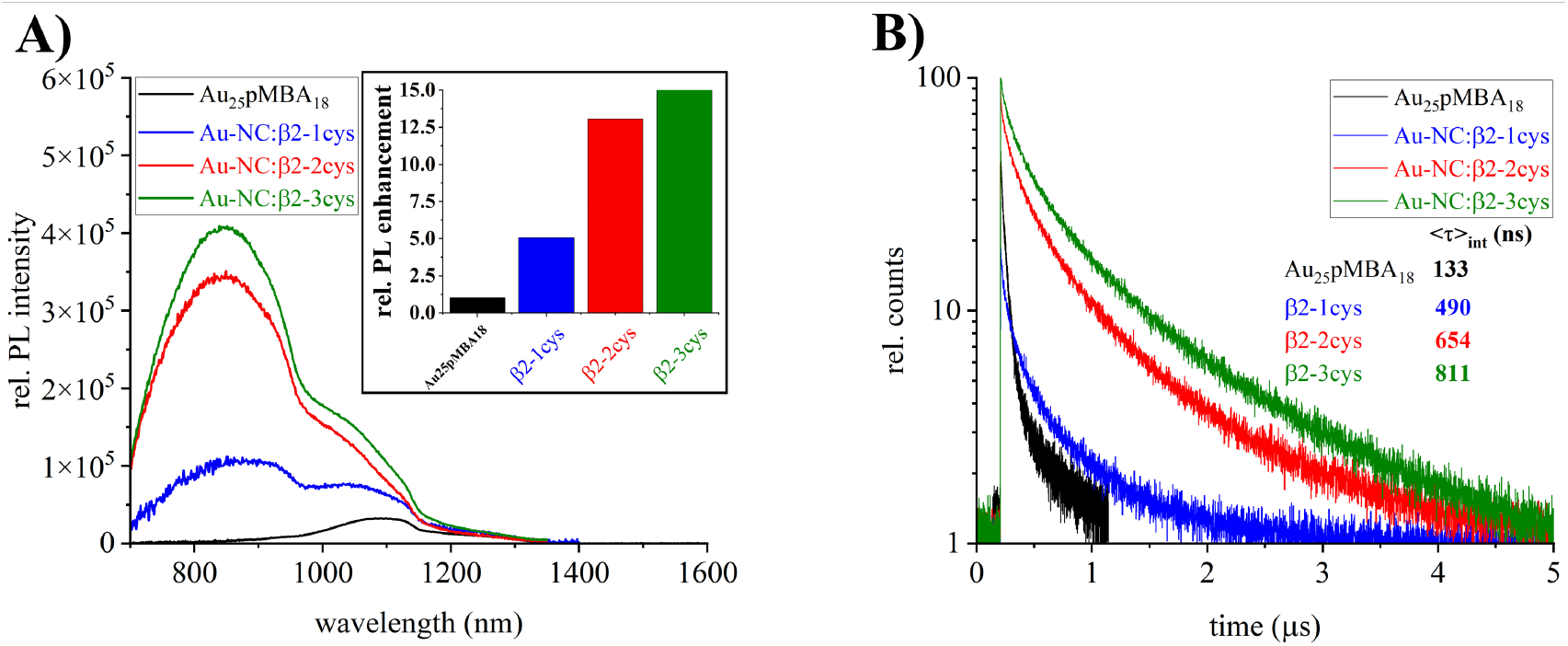
(a) Absorbance corrected PL emission spectra of the major species of the β2 conjugates. λ_exc._ 450 nm. The inset shows the calculated PL enhancement with respect to the isolated Au-NC. (b) PL decay curves of the three major Au-NC:β2 species and the calculated intensity weighted average lifetime. λ_exc._ 510 nm.

The structural characterization of the Au-NC conjugates were performed by Small-Angle X-ray Scattering (SAXS), a biophysical technique that is extremely sensitive to the size and shape of nano-objects in solution^15,16^. Either batch or the Size-Exclusion Chromatography coupled to SAXS (SEC-SAXS) measuring modes were used depending on the amount of conjugate available (Fig. 3A and S6). For the three β2 constructs, the curves obtained for Peak 1 and Peak 2 indicated that these species were very compact, with radii of gyration (*R*_*g*_) between 11.2 and 14.0 Å (Table S2). Note that β2, in the absence of Au-NCs, has a *R*_*g*_ of 24.3 ± 0.1 Å (Fig. 3)^17^. This compact structure can be attributed to the presence of small nanoparticles with a diameter of ∼12 Å with a MW (7.7 kDa) that is similar to that of the protein (8.1 kDa). The presence of dense particles reduces the resulting *R*_*g*_ of the conjugates and strongly perturbs the SAXS curve. Interestingly, when analyzing the dimensions of the major species for the three constructs, where all cysteines were engaged in Au-NC binding, a systematic reduction of the *R*_*g*_ was observed when increasing the number of cysteines (14.0 ± 0.1 Å, 12.2 ± 0.1 Å and 11.3 ± 0.1 Å for β2-1cys, β2-2cys and β2-3cys complexes, respectively). This suggests that the increase in the number of anchoring points reduced the volume of the disordered protein. In agreement with the simultaneous complexation to two proteins, larger sizes were observed for the species corresponding to Peak 3. Again, a compaction was observed when passing from β2-1cys to β2-2cys conjugates (with *R*_*g*_s of 17.1 ± 0.2 Å and 14.9 ± 0.1 Å, respectively) due to the larger number of cysteines bound to the Au-NCs. A different scenario was observed for Peak 3 of the Au-NC:β2-3cys, which has a *R*_*g*_ closer to the β2-1cys (17.5 ± 0.1 Å), probably due to the presence of β2 chains engaging a single cysteine in the complex (Fig. 1). Despite the compact structure of the conjugates, the absence of a maximum in the Kratky representations of all measured SAXS curves indicated that the protein chains displayed an important degree of mobility upon binding to the Au-NC (Fig. 3B and S7). Interestingly, the slope of the Kratky plots at large angles, which also reports on the compactness of the sample, was in line with the number of cysteines of the β2 construct and the *R*_*g*_ values^18,19^.

**Figure 3.**
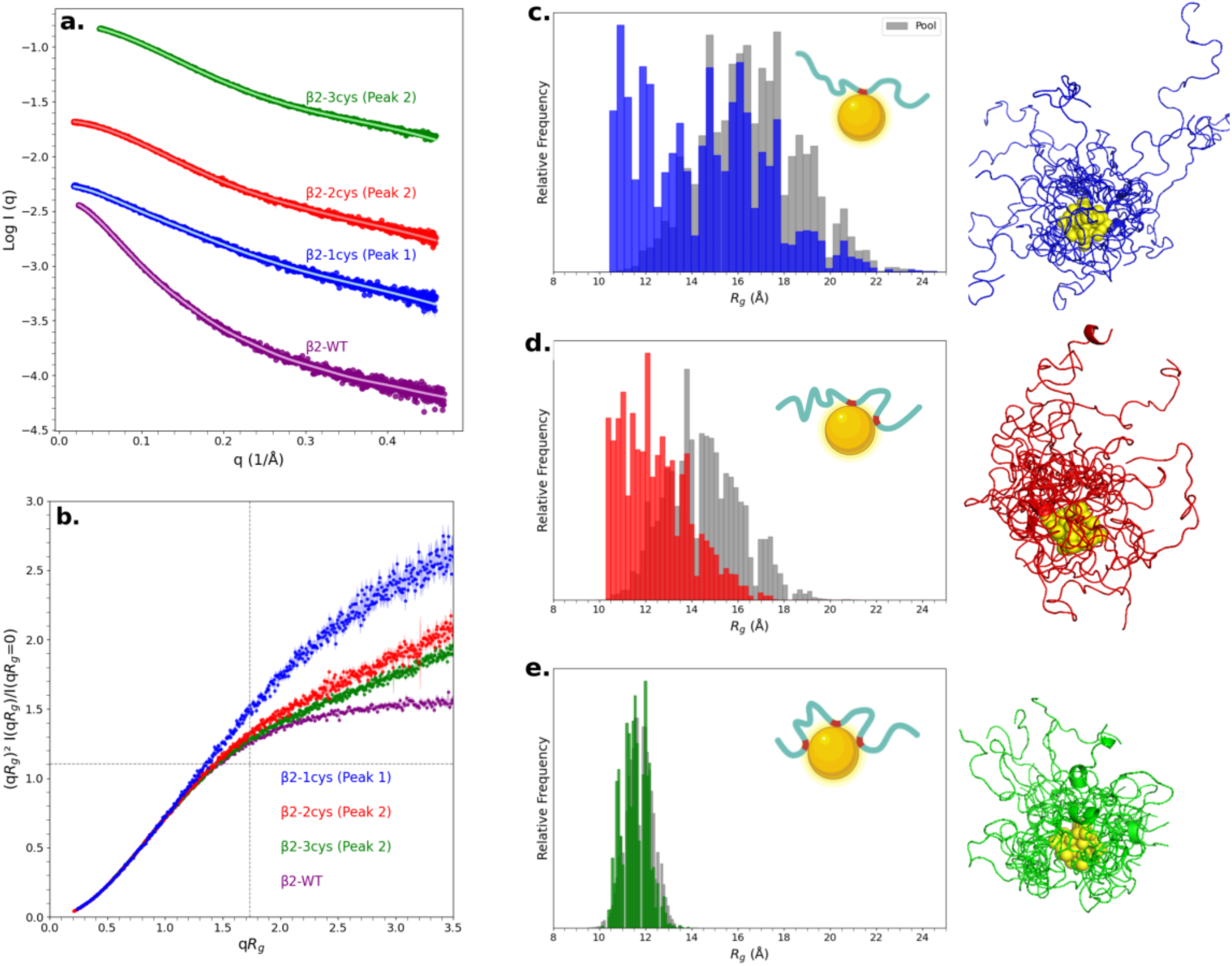
**(a)** SAXS curves for the β2-1cys, β2-2cys, β2-3cys main peaks and β2-WT, which contains two free cysteines. **(b)** Kratky plots for the β2-1cys, β2-2cys, β2-3cys main peaks and β2-WT. **(c)** R_g_ distribution for β2-1cys Peak 1, with its corresponding ensemble. **(d)** R_g_ distribution for β2-2cys Peak 2, with its corresponding ensemble. **(e)** R_g_ distribution for β2-3cys Peak 2, with its corresponding ensemble.

We then used the SAXS data to derive detailed structural models for the different conjugates to analyze the conformational behavior of β2 when bound to an Au-NC. Therefore, we first built atomic models of all assemblies using the MS information regarding the number of cysteines engaged in the complex (Fig. 1). While the coordinates of the Au_25_pMBA_18_ were considered fixed, polypeptide chains were built using sampling methods implemented in the MoMA software suite^20^, placing the bound cysteines at the position of randomly chosen pMBA molecules of the Au-NC (see Method section for details). An excellent description of all SAXS profiles was obtained, with *χ*_2_ values below 1.14, when using the Ensemble Optimization Method (EOM)^21,22^ to fit the data using MS-compatible structural models (Fig. 3A, S6 and Table S2). This indicated that the resulting ensembles were a good representation of the different Au-NC complexes in solution. Sub-ensembles of conformations compatible with the experimental SAXS data of the three major species are displayed in Fig. 3. The comparison between the *R*_*g*_ distributions from the initial pool of conformations and those obtained from the EOM fit is an excellent tool to quantitatively evaluate the compactness of protein ensembles in solution (Fig. 3 and S8)^23,24^. The EOM analyses of the SAXS curves of these major species indicated that the conformational space covered by these conjugates strongly depended on the number of cysteines present in the construct (Fig. 3C-E). The β2-1cys conjugate, which is bound to the Au-NC through a single cysteine, revealed a broad range or *R*_*g*_ values, although the population of very compact conformations, with *R*_*g*_ values smaller than 15 Å, was enormously enriched with respect to these available in the conformational pool. A similar conformational enrichment occurred for the β2-2cys conjugate, even though the available conformational space available for this species was notably reduced due to the double anchoring point. Finally, the conformational space of the β2-3cys conjugate was very narrow, with *R*_*g*_ values from 10 to 14 Å, and very similar to that observed for the available conformations with the three cysteines bound to the Au-NC. These observations suggest that, in addition to the cysteine anchoring, other interactions occur at the interface between the protein and the Au-NC that induce a general increase in particle compactness (Fig. 3C-E). To support this observation, we compared the *R*_*g*_ distributions of the ensemble derived from the EOM fitting for the Peak 2 of the Au-NC:β2-2cys with that of the same ensemble after synthetically removing the Au-NC (Fig. S9). The systematic shift towards smaller *R*_*g*_ values probed the effect of the presence of a heavy Au-NC. When comparing these distributions with the EOM result of β2, which is much more extended, we noticed that the main species of Au-NC:β2-2cys adopted very compact conformations, necessarily originating from the non-covalent interactions of the protein with the Au-NC surface (Fig. S9).

In line with the observations for the major species, the enrichment in very compact IDP conformations was also observed for the minor peaks of the conjugates (Peak 1 of β2-2cys and β2-3cys, and Peak 2 of β2-1cys), suggesting similar interaction mechanisms at the surface of the Au-NC (Fig. S8). A different scenario emerged when analyzing the data corresponding to the Peak 3 for the three β2 constructs, which contain two disordered chains per Au-NC. In this case, the selected conformations displayed size distributions that were very similar to those of the original pools (Fig. S8). We speculate that the steric hindrance between the two proteins hampered extensive non-covalent interaction at the Au-NC interface as occurred with the more compact species containing a single disordered chain.

### Concluding Remarks

In this study, we have elucidated the ensemble structures of distinct bioconjugates formed between an ultrasmall, atomically precise Au-NC and three bioengineered disordered proteins with an increasing number of free cysteines. Through this extensive characterization, we showed that by modifying the stoichiometry, reaction time, and number of anchoring points in the protein, we can precisely control the chemical nature of the resulting species. Moreover, MS data showed that these nanobioconjugates are formed through a ligand exchange mechanism whereby the thiol groups occupy the position of pMBA molecules, removed from the Au-NC surface. These modifications solely occur in the external shell of the Au-NCs, while the internal structure of the metal core is preserved. Our integrative approach indicates that the resulting complexes adopt very compact conformations in solution. While the degree of compactness can be modulated by the number of cysteines of the protein, extensive, nonspecific interactions seem to occur at the interface of the metal cluster and the disordered protein chains, further compacting these complexes. Surprisingly, despite the presence of these covalent and non-covalent interactions at the interface, the protein remained completely disordered, covering the surface of the Au-NCs. Although this study has been performed for an ultrasmall NC containing 25 Au atoms, we have observed that these bioconjugates are also formed in the presence of a larger NC containing 102 Au atoms (see Fig. S10). This observation suggests that the mechanisms unveiled in the present study could be general for pMBA-protected Au-NCs.

These structural changes are accompanied by significant changes in the optical properties of the nanobioconjugates, which depended on the number of cysteine residues present in the proteins. A relatively strong enhancement of the emission band centered around 850 nm (upon excitation at 450 and 690 nm; see Fig. S4) was observed and a modification of the contribution of the different emissive bands to the overall PL with its initial PL maximum at 1100 nm is consistent with previous observations for the attachment of this Au-NC to cysteine-34 of BSA ^8^. However, exchanging pMBA for the electron-rich cysteine residue alone cannot fully account for the observed increase in PL intensity and spectral effects. More likely, the location of the disordered protein chains on the Au-NC surface, reduces PL quenching by water molecules which are excluded from the Au-NC surface. Moreover, the extensive covalent and non-covalent interactions between Au-NC and the protein rigidify molecules at the interface and reduce the ICT, which in turn lead to the overall increase in PL emission and prolonged PL lifetimes.

In summary, this work establishes general design principles to guide the development of nanobioconjugate libraries combining metal nanoclusters and engineered proteins. Through synergistic mechanisms mainly occurring at the interface, these conjugates display properties that are different from those of the individual components and that can be rationally tuned. Our results open new avenues for tailoring nanodevices for biomedical applications, such as selective biosensing, targeted therapeutics, and protein labeling tools that preserve native protein structure.

## ACKNOWLEDGMENT

The CBS is a member of France-BioImaging (FBI) and the French Infrastructure for Integrated Structural Biology (FRISBI), two national infrastructures supported by the French National Research Agency (ANR-10-INBS-04-01 and ANR-10-INBS-05, respectively). The authors acknowledge SOLEIL (Saint-Aubin, France) for the provision of synchrotron radiation facilities (proposal 20220968).

## ABBREVIATIONS

NC: nanoclusters
IDP: intrinsically disordered protein
NIR: near-infrared
pMBA: para-mercaptobenzoic acid
cys: cysteine
*R*_*g*_: gyration radius
PAGE: Polyacrylamide Gel Electrophoresis
MS: Mass Spectrometry
MW: Molecular Weight
PL: photoluminescence
SAXS: Small-Angle X-ray Scattering
SEC: Size-Exclusion Chromatography
EOM: Ensemble Optimization Method
ICT: intramolecular charge transfer
BSA: Bovine Serum Albumin.

