## Supplementary material for "Interplay Between Intrinsically Disordered Proteins and Atomically Precise Gold Nanoclusters Modulates their Optical Properties": supplentary material

**# Equivalent contribution**

### Experimental section

#### Synthesis and purification of atomically precise Au-NCs

The synthesis of $Au_{25}pMBA_{18}$ followed the protocol by Bertorelle *et al.* (1). First, 79 mg of 4-Mercaptobenzoic acid (pMBA) were dissolved in 40 mL of methanol and 4 mL of tributylamine. Then, 100 mg of tetrachloroauric acid trihydrate were added at room temperature and stirred 30 minutes to form a gold thiolate/tributylamine complex. Then, to induce a slow reduction of gold, 200 mg of trimethylamine borane were added under stirring for 2 h before adding 200 mg more. The solution was left under stirring overnight to achieve the formation of the clusters.

The precipitation of the clusters was induced by adding 1 mL of 10% $NH_4OH$ solution and 30 mL of diethyl ether ($Et_2O$). After centrifugation, the supernatant, which contains unwanted products, was removed. A second redispersion/precipitation cycle (1 mL $H_2O$; 5 mL of MeOH and 20 mL of $Et_2O$) was done. Purification of the $Au_{25}pMBA_{18}$ clusters was done by selective precipitation. The precipitate was redispersed in 5 mL of water and then 700 mg of ammonium acetate were added, followed by 20 mL of MeOH. Under these conditions, a precipitate forms and it is separated by centrifugation. The precipitate was discarded and the supernatant was further precipitated by adding $Et_2O$. Two cycles of redispersion/precipitation were done to improve purity of the cluster. The pellet was then resuspended in water and containing 100 mM Triethylammonium acetate to achieve a final concentration of 1 mg/mL.

**Protein constructs**

The C-terminal part of the β2-adrenergic receptor (β2) was cloned into a pET-1a expression vector as described by Guillien et *al.* (2). This plasmid, pETGST-1a-β2AR WT, allowed the expression of a (His)6-GST-tagged protein. In the wild type sequence, the β2 carries two natural cysteines at the positions 39 and 67. Here, we have engineered different mutants named pETGST-1a-β2AR C39A+C67A, pETGST-1a-β2AR C67A and pETGST-1a-β2AR S16C containing zero, one or three cysteines, respectively (Table S3). All these vectors were constructed using the QuikChange Site-directed mutagenesis kit (Agilent™). pETGST-1a-β2AR C67A and pETGST-1a-β2AR S16C plasmids were first synthesized using pETGST-1a-β2AR WT vector as a template, and then pETGST-1a-β2AR C39A + C67A was constructed using vector pETGST-1a-β2AR C67A as a template.

**Protein expression and purification**

All the plasmids were transformed in *Escherichia coli* BL21 DE3 strain (Fisher Scientific, Illkrich, France). Proteins were expressed in self-inducible medium ZYM50-52 at 37°C. After harvesting, bacteria were resuspended and lysed by sonication in 20 mM Tris pH 7.5, 300 mM NaCl, 2 mM dithiothreitol (DTT) named Buffer A and supplemented with anti-protease (Complete EDTA free tablet (Roche, Saint Quentin Fallavier, France)). Cell debris were removed by centrifugation, the soluble fraction was loaded into a 5 mL HisTrap Excel column (Cytiva, Uppsala, Sweden, 17371906). The column was washed in Buffer A supplemented with 10 mM imidazole. Proteins were eluted with a linear gradient of Buffer A supplemented with 500 mM imidazole. The eluate was dialysed in Buffer A containing TEV protease (1/50 (w/w) protease/protein) at 4°C overnight. Proteases and His-GST tag were removed using a gravity column containing Glutathione Sepharose 4B (GE Healthcare, Uppsala, Sweden, 17-0756-05) and Complete His-Tag purification resin (Roche, Mannheim, Germany, 05893682001). Proteins were concentrated using a 3 kDa Amicon concentrator (Merck, Cork, Ireland, UFC900324) then injected into a HiLoad 16/600 Superdex 75pg column (GE Healthcare, Uppsala, Sweden, 28-9893-33) with 50 mM Bis Tris pH 6.7, 50 mM NaCl, 2 mM DTT. The purified proteins were stored at -20°C until use.

**Preparation and purification of the Au-NC:β2 complexes**

The buffer protein was exchanged to PBS-1X supplemented with 100 μM TCEP (Tris(2-CarboxyEthyl)Phosphine) then diluted to 100 μM. The mixing of proteins with Au-NC at a 1:2 ratio (15 μM : 30 μM) was done at room temperature with low agitation for 3h. The mixture was concentrated to 10X using a 3 kDa Amicon concentrator (Merck, Cork, Ireland, UFC800324) then injected into a Superdex 75 10/300 GL column (GE Healthcare, Uppsala, Sweden, 17-5174-01) equilibrated in PBS-1X. Purified samples were stored at -20°C until use.

**Polyacrylamide Gel Electrophoresis (PAGE)**

PAGE separation was carried out using a vertical gel electrophoresis unit with a size of 0.2 cm × 20 cm × 20 cm. The separating and stacking gels were prepared from acrylamide monomers with the total contents of 20 and 7 wt %, respectively [40% acrylamide–bis(acrylamide) 29:1]. The eluting buffer is a standard TBE buffer (89 mM Tris, 89 mM boric acid, 2 mM ethylenediaminetetraacetic acid). The Au-NC:β2-2Cys complexes were dissolved in a 15% (v/v) glycerol/water solution. The sample solutions were loaded onto the stacking gel (200 μL) and eluted for 2 h at a constant voltage mode (150 mV) to achieve sufficient separation of the peaks. The gel was then removed from the casket and placed under either a visible-NIR camera with a couple excitation/emission at 647nm/667 nm (Typhoon FLA 9500 (GE Healthcare Life Sciences)) or under a SWIR camera (Nirvana 640-ST coupled with 35 mm lens (Navitar) and long pass filter at 900 nm (Thorlabs) following an excitation with a Laser at $\lambda_{exc.}$= 808 nm.

**HPLC-coupled mass spectrometry**

Purified samples were concentrated to at least 10X using a 3 kDa Amicon concentrator (Merck, Cork, Ireland, UFC800324) before the Mass spectrometry (MS) analyses. MS was performed at the OLAF Mass spectroscopy platform of the University of Montpellier with a Synapt G2-S (Waters, UEB205) equipped with an electrospray ionization source. The spectrometer was connected to a UPLC Acquity H-Class (Waters) equipped with a Bioresolve RP mAb 150x2.1mm, 1.7 µm (Waters) column and the UV chromatogram was recorded at 280 nm. Data were recorded using positive and negative ionization modes, although only results obtained using the positive mode are reported.

**Optical and spectroscopic characterization of complexes**

Absorbance measurements were performed using a Cary5000 (Agilent) UV-Vis-NIR spectrophotometer in the range of 300-1100 nm. Steady-state photoluminescence spectra were measured from 700–1600 nm with a calibrated FSP 920 (Edinburgh Instruments, Edinburgh, United Kingdom) spectrofluorometer equipped with a nitrogen-cooled PMT R5509P. Time-resolved measurements were performed in the wavelength region of 900 - 940 nm using a FLS 920 (Edinburgh Instruments, Edinburgh, United Kingdom) lifetime spectrofluorometer equipped with an EPL-510 (Edinburgh Instruments, Edinburgh, United Kingdom) picosecond pulsed diode laser (excitation wavelength of 510±10 nm; power of 5 mW) and a fast PMT R2658P from Hamamatsu, respectively. The PL decays were fitted with a multi-exponential model, see Eq. (1)

$$I(t) = \sum_{i}^{n} A_i \; e^{\left(\frac{-t}{\tau_i}\right)} \qquad (1)$$

where $A_i$ are the amplitudes and τ the lifetimes of the respective decaying species. These data were calculated with a reconvolution fitting procedure using the FLS-920 spectrometer software. The fit quality was evaluated from the corresponding $\chi^2$ values. The average lifetimes were calculated as an intensity-weighted average, see Eq. (2)

$$< \tau >_{int} = \frac{\sum_{i}^{n} A_i \tau_i^2}{\sum_{i}^{n} A_i \tau_i} \qquad (2)$$

where, $\tau_i$ are the lifetimes of each component and $A_i$ the corresponding relative fractional amplitudes.

**Small-angle X-ray scattering measurement and raw data analyses**

Small-angle X-ray scattering (SAXS) data were acquired for the different purified peaks for the Au-NC:β2 conjugates at the SWING beamline at the SOLEIL Synchrotron (Saint-Aubin, France) (3) using X-ray wavelength of 1.03 Å and a sample-to-detector distance of 2.00 m. Samples were measured at 15°C either in batch or in Size-Exclusion Chromatography coupled to SAXS (SEC-SAXS) modes in PBS-1X. For the SEC-SAXS mode, 45 µL of samples were injected into a 3 mL Superdex 75 5/150 GL column (Cytiva, Uppsala, Sweden) pre-equilibrated into the same buffer and were eluted at 0.2 mL/min.

The scattering profiles measured covered a momentum transfer range of 0.002 $Å^{-1} < q < 0.5$ $Å^{-1}$. In SEC-SAXS experiments, the scattering patterns of the buffer were recorded before the void volume of the column (1 mL). SAXS data were processed using CHROMIX (4) to automatically select frames corresponding to buffer and sample, perform buffer subtraction and average profiles. The averaged SAXS curves were analyzed using Primus from ATSAS software package (5).

#### SAXS-driven modelling of the gold-conjugated nanostructures

The modeling of each of the Au-NC:β2 complexes was performed using the conformational sampling algorithms implemented in the MoMA software suite (https://moma.laas.fr/) (6). For species with a single cysteine engaged in the interaction (Peak 1 and Peak 2 of Au-NC:β2-1Cys, and Peak 1 of Au-NC:β2-3Cys), the building procedure was the following: (*i*) A single extended conformation of β2 was collected and an Au-NC particle with 17 pMBA molecules was manually approached with Pymol to the free cysteine, positioning the free gold atom (after removing the pMBA) at 2.2–2.6 Å to the sulfur atom (7). (*ii*) From this initial state, a conformational ensemble (20,000 conformations) of β2 was built with MoMA (6) by fixing the position of the bound cysteine. (*iii*) The sampled conformations were submitted to a filtering procedure in order to select those sterically compatible with the Au-NC. Concretely, protein conformations with at least one atom within 1 Å of any gold atom were discarded, unless that atom was in the flanking region (±3 residues) of the bound cysteine. A similar procedure was used to build species with 2 or 3 anchored cysteines and with two different chains. The main difference was the initial β2 conformations, which were built by forcing MoMA to position the anchored cysteines in close proximity to make chemically compatible interactions with the Au-NC after removing the specified number of pMBA molecules. In all the cases, the filtering criteria were maintained.

After filtering, the SAXS profile for each conformation of each conjugate was calculated using CRYSOL (8), setting the number of spherical harmonics to 35 and treating hydrogen atoms explicitly. The calculated SAXS profiles were fitted to experimental data using GAJOE from the

EOM suite (9,10), running 500 cycles of 2,000 generations each, with an ensemble size of 10, disallowing repetitions.

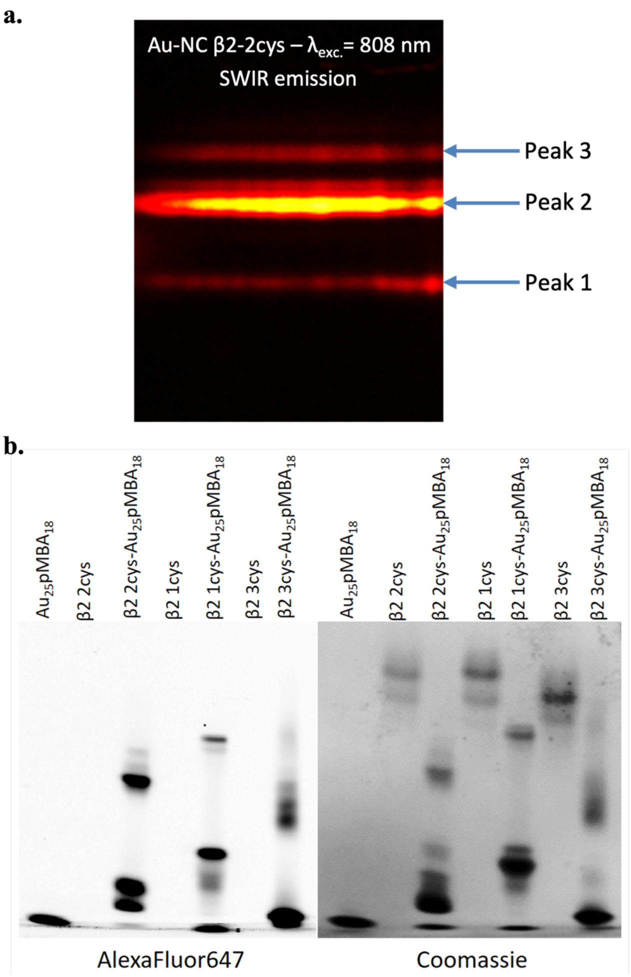


**Figure S1. (a)** Photoluminescent image of PAGE migration profile of Au-NC: β2-2cys complexes in a native gel in the SWIR following an excitation at $\lambda_{exc.}$= 808 nm. **(b)** Native PAGE-gel revealed by fluorescence emission at 667 nm (left) and Coomassie (right) of the mixtures of the $Au_{25}pMBA_{18}$ with the 3 β2 constructs.

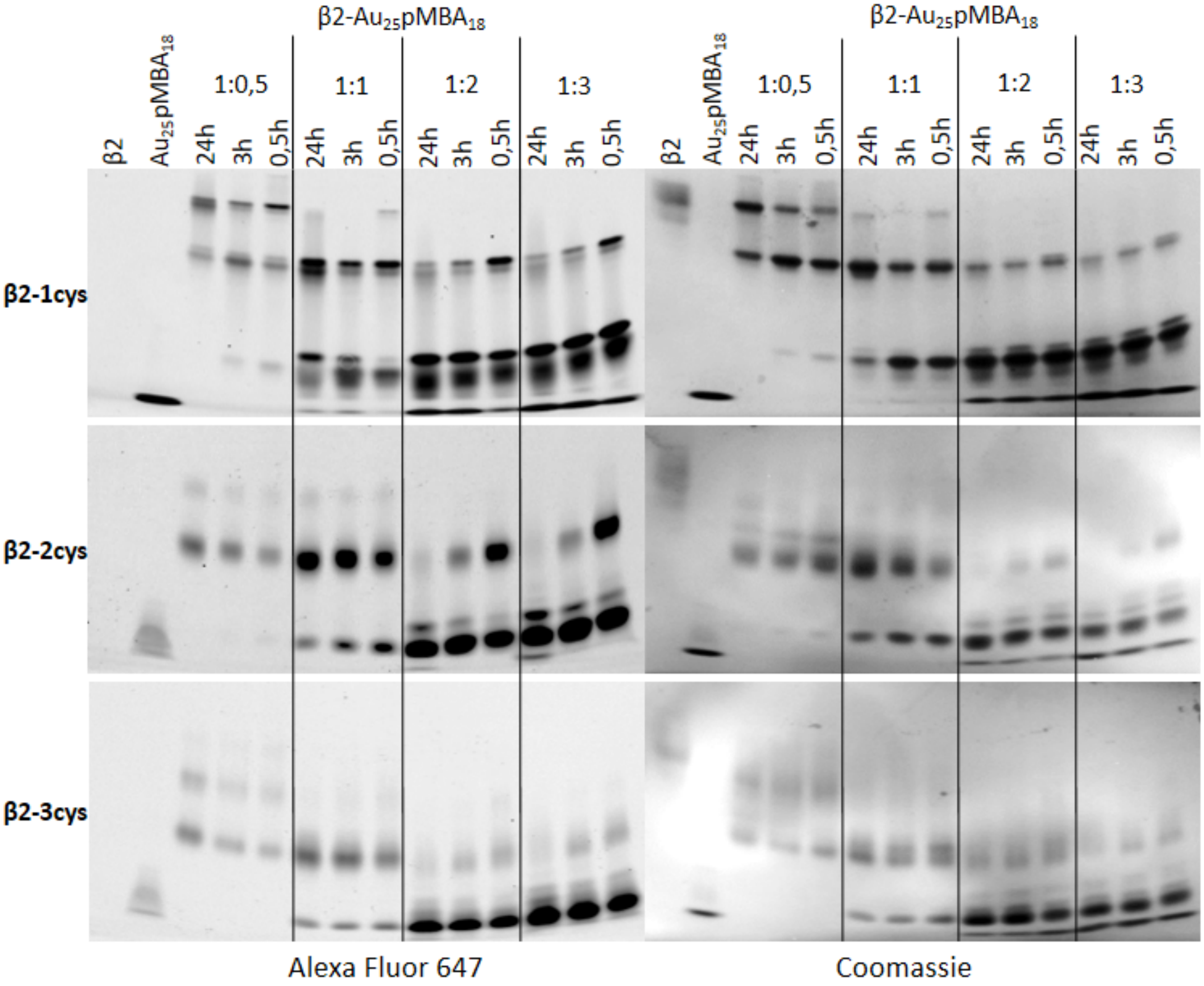


**Figure S2.** Population of the bioconjugates formed upon reacting $Au_{25}pMBA_{18}$ with the 3 β2 constructs (β2-1cys, β2-2cys and β2-3cys from top to bottom) depending on the reaction time (24h, 3h and 0.5h) and the protein:Au-NC stoichiometry (1:0.5, 1:1, 1:2, 1:3). Native PAGE-gels have been revealed by fluorescence emission at 667 nm (left) and Coomassie (right).

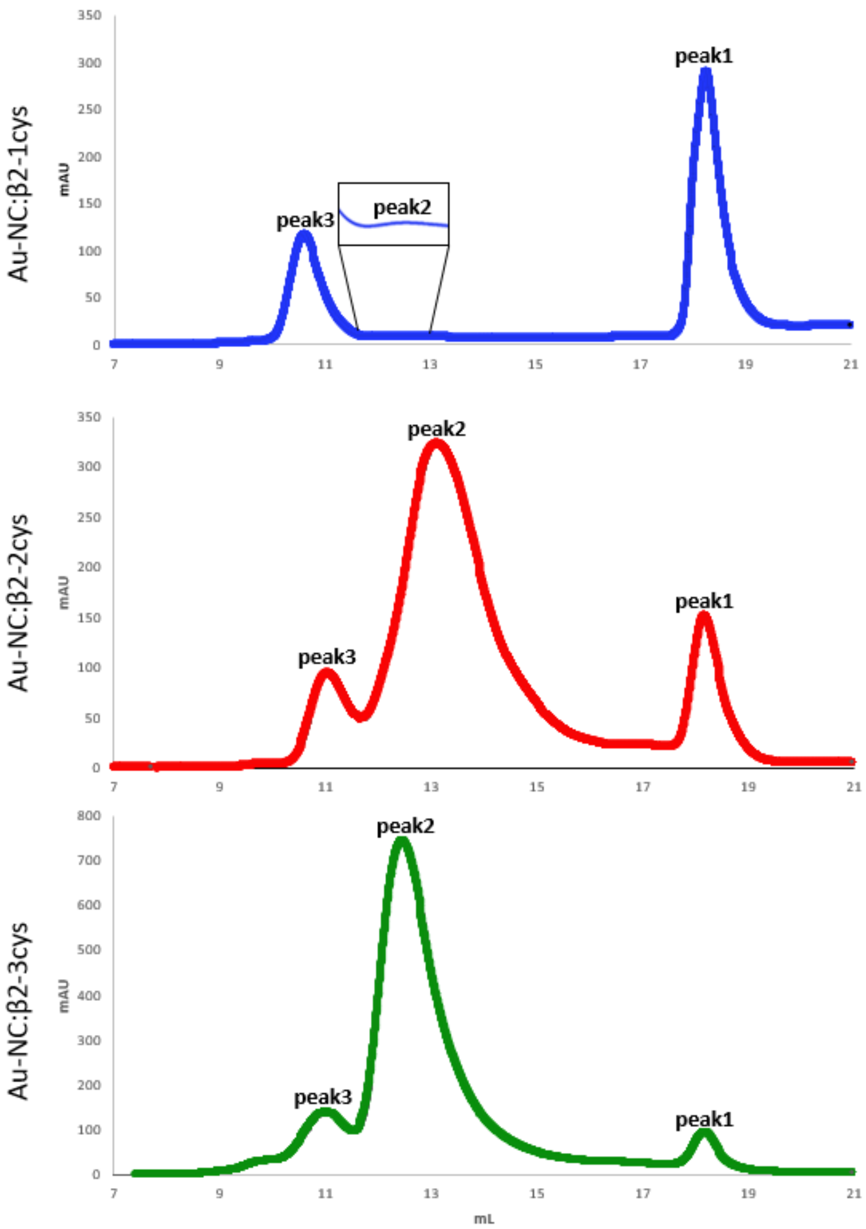


**Figure S3.** UV profiles of the Size-Exclusion Chromatography obtained after mixing $Au_{25}pMBA_{18}$ with β2-1cys (blue), β2-2cys (red) and β2-3cys (green). The individual fractions were collected and concentrated for the subsequent optical and structural characterization.

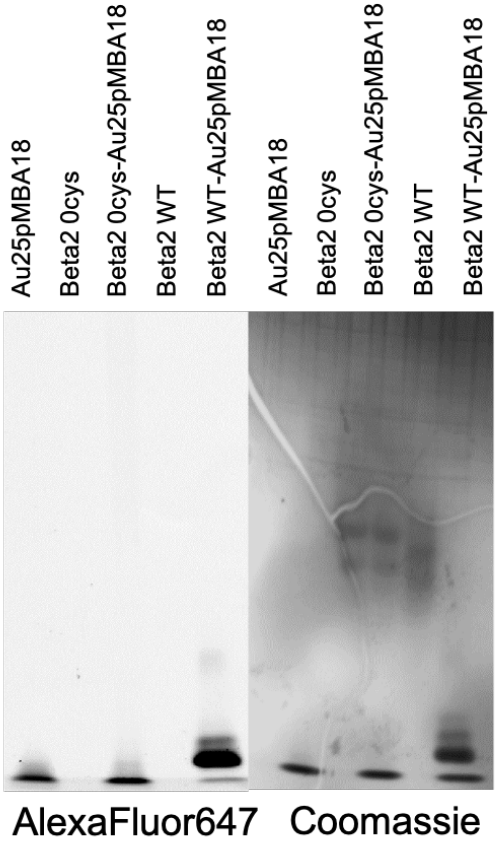


**Figure S4.** Native PAGE-gel revealed by fluorescence emission at 667 nm (left) and Coomassie (right) of the mixtures of the $Au_{25}pMBA_{18}$ with the β2-0cys construct. The gel shows that no complex is formed when mixing the Au-NC with a protein without free cysteines.

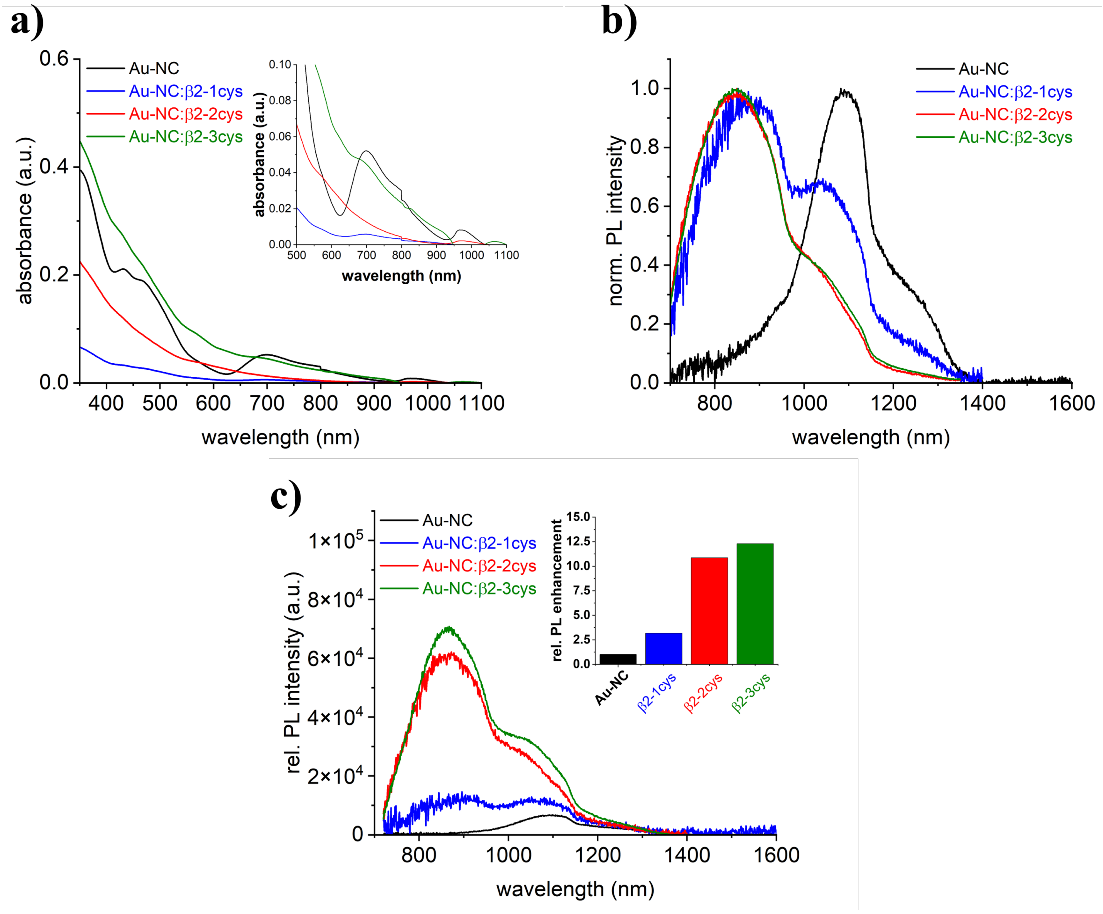


| Sample | $A_1$ | $\tau_1$ (ns) | $A_2$ | $\tau_2$ (ns) | $A_3$ | $\tau_3$ (ns) | $<\tau>_{int}$ (ns) |
|---|---|---|---|---|---|---|---|
| $Au_{25}pMBA_{18}$ | 0.040 | 6.0 | 0.122 | 44.1 | 0.016 | 259.4 | **133** |
| Au-NC:β2 1-cys | 0.171 | 22.5 | 0.173 | 153.9 | 0.083 | 680.0 | **490** |
| Au-NC:β2 2-cys | 0.143 | 35.3 | 0.184 | 250.9 | 0.095 | 908.2 | **654** |
| Au-NC:β2 3-cys | 0.125 | 48.1 | 0.172 | 278.7 | 0.110 | 1068.1 | **811** |

**Figure S5. (Top) A)** Absorbance spectra of the major species of the β2 conjugates with an enlarged representation of the spectra in the range from 500 to 1100 nm as inset. **B)** Normalized PL spectra of the major species of the β2 conjugates excited at 450 nm. **C)** Absorbance corrected PL emission spectra of the major species of the β2 conjugates excited at 690 nm. The inset shows the calculated PL enhancement. **(Bottom)** Summary of the amplitudes ($A_i$) and PL lifetimes ($\tau_i$) obtained from fitting the PL decay curves using a three exponential fit and the calculated intensity weighted average PL lifetime ($<\tau>_{int}$).

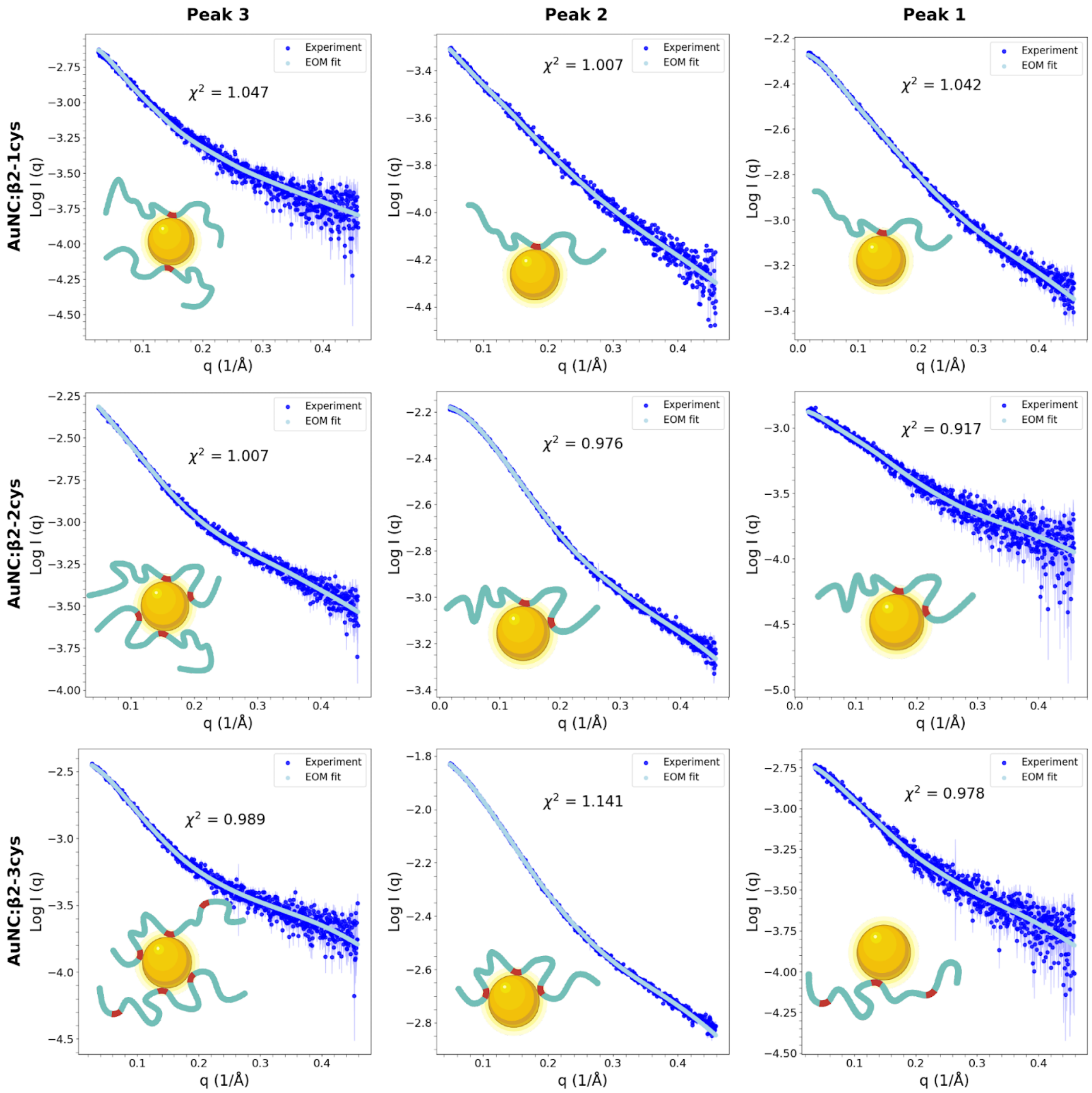


**Figure S6.** SAXS intensity profiles (blue dots) as a function of the momentum transfer, $q$, for each of the peaks purified after mixing Au-NC with the three β2 constructs. The fit using the Ensemble Optimization Method (EOM) is displayed as a solid cyan line. Cartoons displaying the topology for each species obtained by mass spectrometry and modelled for the subsequent fitting are displayed in the panels.

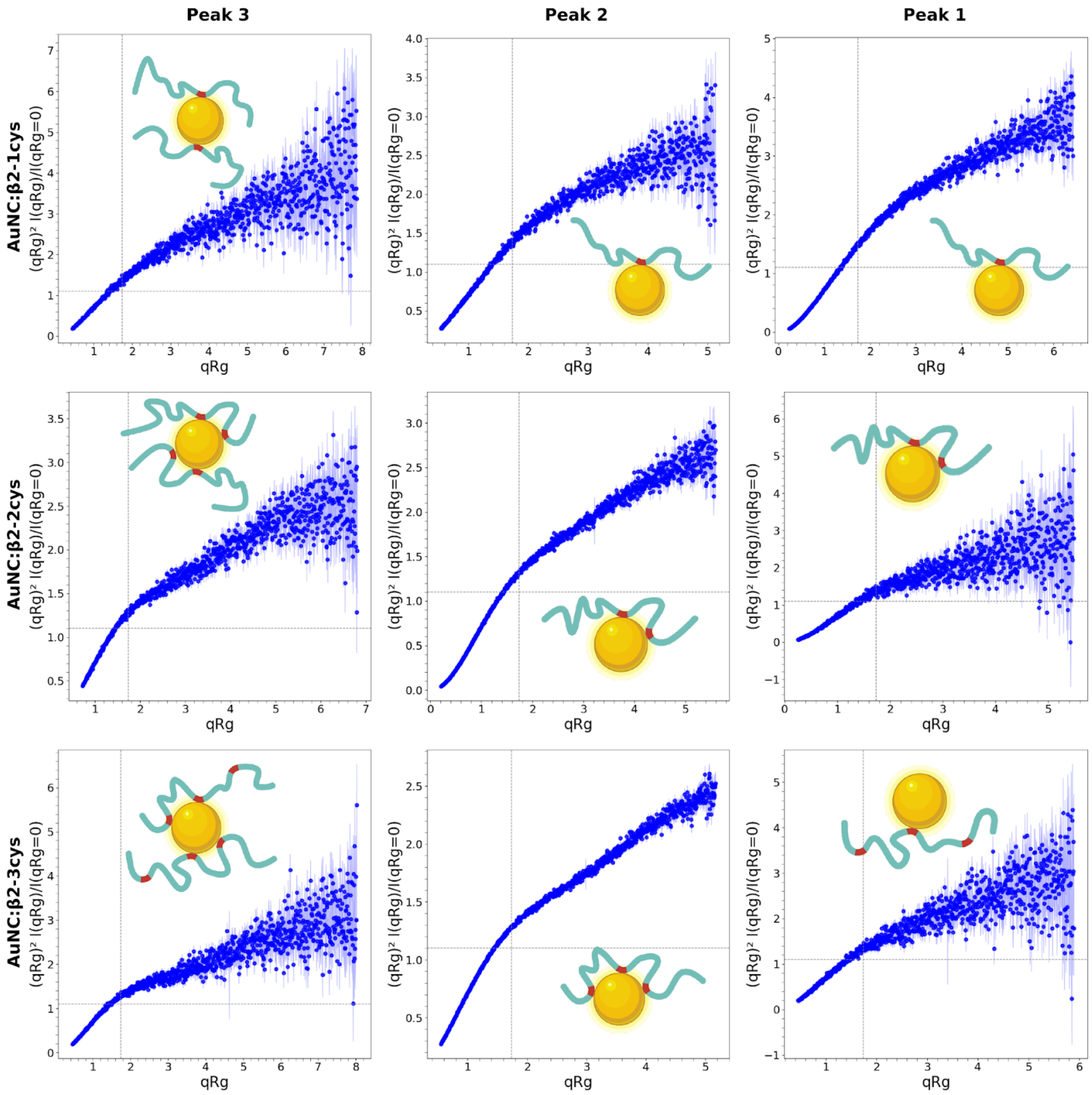


**Figure S7.** Dimensionless Kratky plots derived from the SAXS profiles (blue dots) for each of the peaks purified after mixing Au-NC with the three β2 constructs. Cartoons displaying the topology for each species obtained by mass spectrometry and modelled for the subsequent fitting are displayed in the panels.

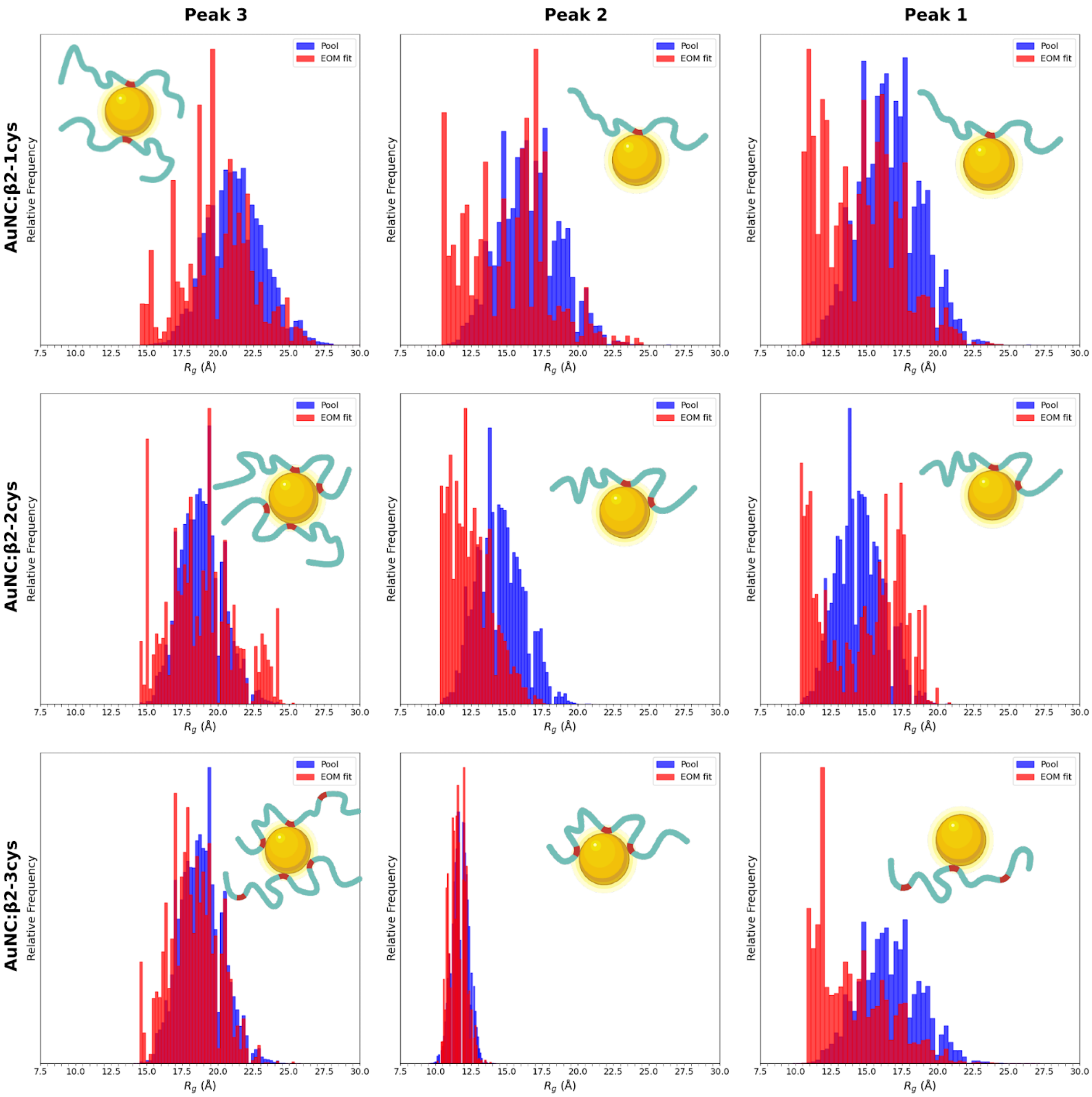


**Figure S8** Comparison of the $R_g$ distributions from the pool of conformations (blue) with these derived from the EOM analysis (red) of the SAXS curves measured for each of the peaks purified after mixing Au-NC with the three β2 constructs. Cartoons displaying the topology for each species obtained by mass spectrometry and modelled for the EOM fitting are displayed in the panels.

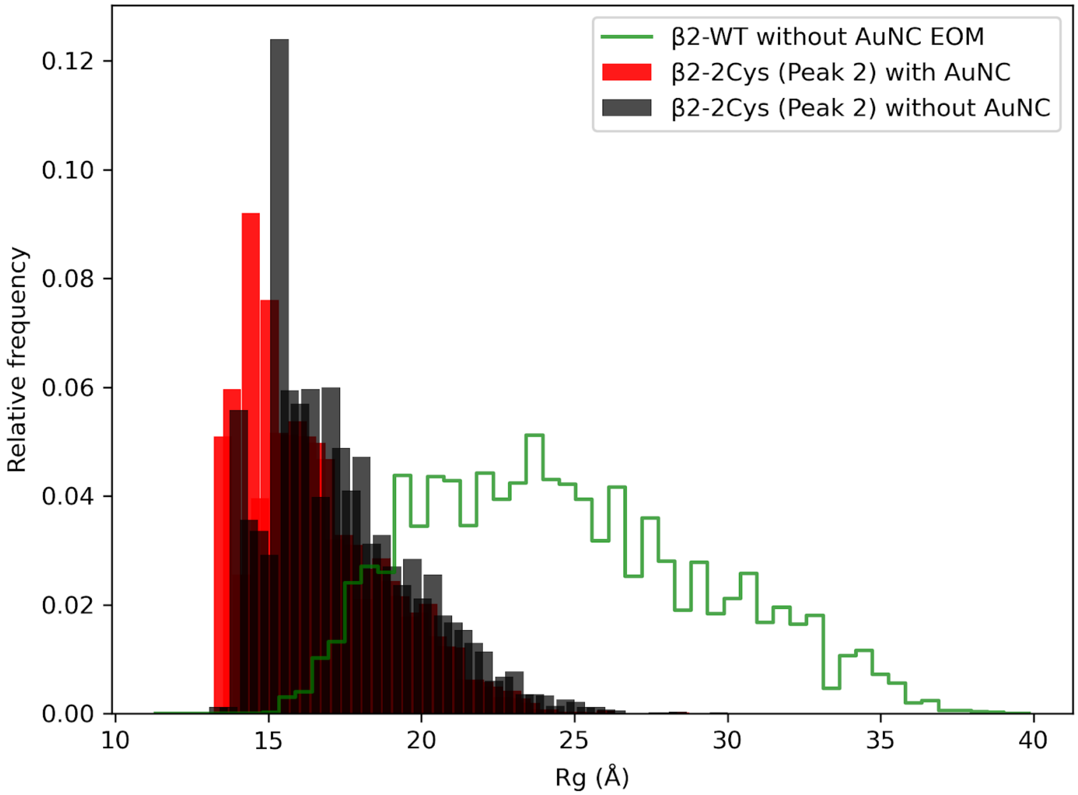


**Figure S9.** $R_g$ distribution corresponding to the EOM analysis of the Peak 2 of the β2-2cys (red), compared to the $R_g$ distribution of the same ensemble in which the Au-NC has been computationally removed before the CRYSOL calculation (gray). In green, the $R_g$ distribution derived from the EOM analysis of the protein β2. These data indicate that, in the presence of the Au-NCs, the protein displays very compact conformations, and that the compactness mainly arises from the non-covalent interactions between the protein and the nanoparticle.

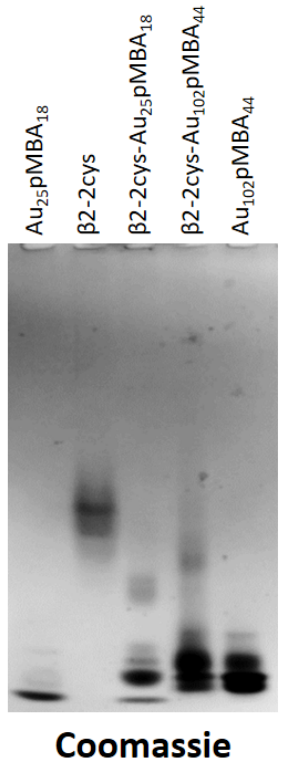


**Figure S10.** Generalization of the ligand exchange mechanism to other Au-NCs. Native PAGE gel of the mixtures of $Au_{25}pMBA_{18}$ and $Au_{102}pMBA_{44}$ with β2-2cys. In both cases additional bands, different from the free protein and the Au-NC, appear in the gel indicating the complexation between both molecules.

**Table S1 :** Amino acid sequences of the β2 constructs used. Cysteines are highlighted in red.

| Protein | Mutations | Sequence |
|---|---|---|
| **β2-0cys** | C39A+C67A | GAMALRRSSLKAYGNGYSSNGNTGEQSGYHVEQEKENKLLAEDLP GTEDFVGHQGTVPSDNIDSQGRNASTNDSLL |
| **β2-1cys** | C67A | GAMALRRSSLKAYGNGYSSNGNTGEQSGYHVEQEKENKLLCEDLP GTEDFVGHQGTVPSDNIDSQGRNASTNDSLL |
| **β2-2cys** | WT | GAMALRRSSLKAYGNGYSSNGNTGEQSGYHVEQEKENKLLCEDLP GTEDFVGHQGTVPSDNIDSQGRNCSTNDSLL |
| **β2-3cys** | S16C | GAMALRRSSLKAYGNGYCSNGNTGEQSGYHVEQEKENKLLCEDLP GTEDFVGHQGTVPSDNIDSQGRNCSTNDSLL |

**Table S1**: Mass spectrometry and SAXS analyses of the purified Au-NC:β2 conjugates

| | **β2-1cys** | | | | | | **β2-2cys** | | | | | | **β2-3cys** | | | | | |
|---|---|---|---|---|---|---|---|---|---|---|---|---|---|---|---|---|---|---|
| **$MW_{Theo}$ (Da)** | 8117.7 | | | | | | 8149.7 | | | | | | 8165.8 | | | | | |
| | $MW_{Exp}$ (Da) | Comp. β2/pMBA | $MW_{Theo}$ (Da) | *SAXS Mode* | $R_g$ (Å) | $\chi^2$ | $MW_{Exp}$ (Da) | Comp. β2/pMBA | $MW_{Theo}$ (Da) | *SAXS Mode* | $R_g$ (Å) | $\chi^2$ | $MW_{Exp}$ (Da) | Comp. β2/pMBA | $MW_{Theo}$ (Da) | *SAXS Mode* | $R_g$ (Å) | $\chi^2$ |
| **Peak 1** | 15644 | 1/17 | 15643.7 | BATCH | 14.0 +/- 0.1 | 1.04 | 15540 | 1/16 | 15522.7 | BATCH | 12.0 +/- 0.1 | 0.92 | 15691.8 | 1/17 | 15691.8 | SEC | 12.9 +/- 0.1 | 0.98 |
| **Peak 2** | 15735 | 1/17 | 15643.7 | BATCH | 11.2 +/- 0.1 | 1.00 | 15515 | 1/16 | 15522.7 | SEC | 12.2 +/- 0.1 | 0.98 | 15389.0 | 1/15 | 15385.8 | SEC | 11.3 +/- 0.1 | 1.14 |
| **Peak 3** | 23606 | 2/16 | 23608.3 | BATCH | 17.1 +/- 0.2 | 1.05 | 23382 | 2/14 | 23366.5 | BATCH | 14.9 +/- 0.1 | 1.00 | 23412.0 | 2/14 | 23398.6 | SEC | 17.5 +/- 0.1 | 0.99 |

**MW:** Au 197 Da, pMBA 153 Da, $Au_{25}pMBA_{18}$ 7679 Da.

**Table S2:** Design of β2 constructs

| Protein | Mutation | Primers | sequence 5'-3' |
|---|---|---|---|
| **β2-0cys** | C39A+C67A | Oligo MUT b2AR 1 Cys | AAAGAAAACAAACTGCTGgcCGAAGATCTGCCTGGCACC |
| | | Oligo MUT b2AR 1 Cys C | GGTGCCAGGCAGATCTTCGgcCAGCAGTTTGTTTTCTTT |
| | | Oligo MUT b2AR 2 Cys | GATAGCCAGGGTCGTAATgcTAGCACCAATGATAGCCTG |
| | | Oligo MUT b2AR 2 Cys C | CAGGCTATCATTGGTGCTAgcATTACGACCCTGGCTATC |
| **β2-1cys** | C67A | Oligo MUT b2AR 2 Cys | GATAGCCAGGGTCGTAATgcTAGCACCAATGATAGCCTG |
| | | Oligo MUT b2AR 2 Cys C | CAGGCTATCATTGGTGCTAgcATTACGACCCTGGCTATC |
| **β2-3cys** | S16C | Oligo b2AR S16C 5’ | CATATGGTAATGGTTATtgcAGCAATGGCAATACCGG |
| | | Oligo b2AR S16C 3’ | CCGGTATTGCCATTGCTgcaATAACCATTACCATATG |
